# Pan-genomic Matching Statistics for Targeted Nanopore Sequencing

**DOI:** 10.1101/2021.03.23.436610

**Authors:** Omar Ahmed, Massimiliano Rossi, Sam Kovaka, Michael C. Schatz, Travis Gagie, Christina Boucher, Ben Langmead

## Abstract

Nanopore sequencing is an increasingly powerful tool for genomics. Recently, computational advances have allowed nanopores to sequence in a targeted fashion; as the sequencer emits data, software can analyze the data in real time and signal the sequencer to eject “non-target” DNA molecules. We present a novel method called SPUMONI, which enables rapid and accurate targeted sequencing with the help of efficient pangenome indexes. SPUMONI uses a compressed index to rapidly generate exact or approximate matching statistics (half-maximal exact matches) in a streaming fashion. When used to target a specific strain in a mock community, SPUMONI has similar accuracy as minimap2 when both are run against an index containing many strains per species. However SPUMONI is 12 times faster than minimap2. SPUMONI’s index and peak memory footprint are also 15 to 4 times smaller than minimap2, respectively. These improvements become even more pronounced with even larger reference databases; SPUMONI’s index size scales sublinearly with the number of reference genomes included. This could enable accurate targeted sequencing even in the case where the targeted strains have not necessarily been sequenced or assembled previously. SPUMONI is open source software available from https://github.com/oma219/spumoni.

## 1 Introduction

Nanopore sequencing instruments have steadily improved in usability, speed, and accuracy. While it lags sequencing-by-synthesis instruments on base quality, quality has improved steadily, with recent datasets reaching and exceeding 90% accuracy [1]. Nanopore sequencing is also convenient and flexible; nanopores are readily used outside of laboratories, e.g. for analyzing biological species in a human or natural environment with the goal of detecting pathogens or contaminants. They can also be used for several assays, including DNA sequencing, direct RNA-sequencing, and the detection of a variety of epigenetic modifications.

Recent computational approaches focus on the problem of allowing nanopores to sequence in a targeted fashion. Oxford Nanopore instruments provide the “Read Until” interface, enabling two-way communication between the sequencer and the control software. The sequencer reports batches of sequencing data, which software can analyze in real time. Importantly, nanopore sequencing has the unique capability where the control software can potentially signal to the sequencer that it should eject the DNA molecule currently in a pore. To eject, the sequencer reverses the voltage across the pore, causing the molecule to reverse direction and exit. The pore is then free to sequence a new molecule. Many such pores – up to 512 per MinION flowcell – are in simultaneous operation; the system can sequence in a targeted manner only as long as the software making ejection decisions can keep up with the aggregate rate of sequencing.

Recently, Payne et al. described the Readfish system [2] which combines an existing base caller with the minimap2 read aligner [3] to align reads to a reference genome in real time and make decisions on whether to eject. The UNCALLED method [4] is similar, but capable of handing the nanopore current signal directly, without first using a base caller. Unlike Readfish, which generally uses a GPU for base calling, UNCALLED is designed to run on a general-purpose CPU. UNCALLED starts by processing the signal to find potential seeds, then maps them to a reference using an FM-index. Finally, it clusters the seeds to identify significant alignments. UNCALLED’s performance degrades as the reference is repetitive, e.g. if it is a collection of related strains.

Motivated by a need for faster methods which can classify reads against large, repetitive references, we developed SPUMONI. For example, in a typical metagenomics experiment, the exact strain or sub-strain of a microorganism is unknown prior to sequencing and therefore, for optimal targeted sequencing, all strains and substrains need to be incorporated into the reference for identification. SPUMONI takes advantage of the overall repetitiveness of these references by building an *r*-index [5], and using the MONI algorithm to calculate matching statistics (MSs) [6].

The *r*-index enables efficient indexing of repetitive collections of reference genomes – e.g. all of the strains of a bacterial species or several human genome assemblies – while still supporting efficient queries. Importantly, the space required by an *r*-index is proportional to the number of runs in the Burrows-Wheeler Transform of the reference genomes (defined as *r*) rather than the total length of the reference genomes. When the collection is highly repetitive, *r* grows sublinearly, and far more slowly than the total length [5].

MONI augmented the *r*-index with an auxiliary data structure enabling more rapid calculation of MSs. An MS at position *i* of a query sequence *P* of length *m* equals the length of the longest prefix of *P*[*i*..*m*] that exactly matches a sequence in the index. MONI efficiently calculates MSs at every position of a query *P*. The first insight of SPUMONI is that these statistics can be used to classify the query sequence; longer MSs indicate a better approximate match to the index.

SPUMONI extends MONI to improve its speed while also making it applicable to the problem of making fast ejection decisions. First, SPUMONI adds a “null index” together with a hypothesis testing framework to make principled ejection choices depending on whether the observed MS lengths are longer than what would be expected by random chance. Second, SPUMONI replaces MONI’s “batch” MS-finding algorithm with a faster online algorithm that calculates a different quantity related to the MS, called the “pseudo matching length”, which we denote as PML (defined in Methods). (SPUMONI stands for Streaming PseUdo MONI.) This optimized PML-finding procedure makes SPUMONI about 3 times faster than MONI, while achieving similar (often greater) accuracy and allowing it to operate on streaming data.

Compared to a minimap2-based approach, SPUMONI can make ejection decisions with respect to a pan-genome index more efficiently. When used to eject bacterial strains in a mock community scenario, SPUMONI has similar accuracy as minimap2 but is about 12 times faster. Moreover, its many-strain index is about 1/15th the size of minimap2’s, and its memory footprint is less than 1/4th the size of minimap2’s. When used to eject simulated human reads in a human microbiome scenario, SPUMONI is faster than minimap2 when both use an index consisting of 3 high-quality human reference genomes. In this scenario, SPUMONI’s memory footprint and index size are higher, although the sublinear scaling of the *r*-index strategy underlying SPUMONI suggests it will benefit from indexes containing many human genomes.

## 2 Results

### 2.1 Method Overview

SPUMONI’s core insight is that a read’s matching statistics (MSs) with respect to an index can reveal whether it has a “good” (i.e. long, high identity) approximate match to the index, without having to perform a more costly read alignment. To determine whether the MSs are long enough to indicate an approximate match, SPUMONI compares the observed distribution of MSs – calculated with respect to a “positive index” containing the target sequences – with those obtained from a “null index” containing the reverse (not the reverse complement) of the sequences from the positive index. The reverse sequences serve as a random sequence of the same length as the positive index but where nucleotide frequencies and simple repeat structures like homopolymers are preserved. As soon as SPUMONI can confidently determine the distributions of MSs from the positive and null indexes are different – possibly having seen only a prefix of the read’s full sequence – it can conclude that the read is among the targets in the positive index. SPUMONI uses a Kolmogorov-Smirnov statistic (KS-stat) threshold to make this decision.

By default, SPUMONI does not generate true matching statistics but instead generates an approximation thereof called Pseudo Matching Lengths (PMLs). These are described in more detail in Methods. SPUMONI can also generate MSs, which it does in its SPUMONI-ms mode.

### 2.2 Experimental Setup

During Nanopore sequencing, electrical current data is transmitted from the sequencing instrument to the control software in “chunks”, representing about 0.4 seconds of sequencing (the exact duration is user-defined parameter). As DNA translocates through the pore at about 450 bases per second, each chunk represents about 180 bases of data. Our experiments on both simulated and real reads mimic the situation where we are processing the first 4 chunks of data delivered by the Read Until API. We chose this time interval as previous work showed it leads to most reads being mapped using minimap2 [2, 3]. We further assume that the data was already base-called, similar to a previous study [2]. In practice, the Read Until API delivers batches of current signal, not bases; we address this further in the Discussion. We did not compare our method to UNCALLED [4] as it is reportedly slower than minimap2 for large genomes and it starts by processing the current signal, where as we have assumed here that we are given base-calls.

With each new batch, both SPUMONI and minimap2 [3] attempt to classify whether the read has an approximate match to a sequence in the positive index. Importantly, SPUMONI deals with new batches of data in an “online” fashion. That is, SPUMONI can easily suspend and resume its MS/PML computation as it awaits a new batch. This is in contrast to minimap2, which takes full reads as input so as to perform full read alignments. Because of this, our evaluation strategy was to run SPUMONI on each batch separately, allowing SPUMONI to possibly make an ejection decision at the end of each of the four batches. Whereas for minimap2, we re-ran minimap2 on successively longer prefixes of the read as new 180-base batches arrived.

After processing a batch, SPUMONI and minimap2 each apply a threshold to determine if the read matches the positive index with high confidence. In practice, this leads to a decision about whether to eject the read. If the positive index contains depletion targets, a high-confidence match to the positive index indicates the read should be ejected. If the positive index contains enrichment targets, the absence of a high-confidence match after some prescribed period indicates the read should be ejected. For our experiments, the positive index always contains depletion (rather than enrichment) targets. Once a method has decided to eject the read, we cease delivering batches for that read; each method is benchmarked only on the read prefix up to the ejection decision, or up to 1.6 seconds (720 bases), whichever comes first.

For the minimap2-based approach, we used the standard ONT settings of minimap2 [3] to align the reads, which are the same settings used by Readfish [2]. We used a MAPQ threshold to decide whether a read was confidently mapped or not. For non-repetitive (“genomic”) references, we used a MAPQ value of 30 or greater to determine if reads were uniquely mapped or not. For repetitive (“pan-genomic”) references, we further checked whether all alignments were to the same species. For further details on the thresholds used, see Section 4.

For evaluation, an instance where a method ejected a read from a genome that was present in the positive index was called a true positive (TP). An instance where a method ejected a read that was not in the positive index was called a false positive (FP). An instance where a method failed to eject a read that was from a positive-index genome was called a false negative (FN).

We performed all the experiments on a computer with a 2.0 GHz Intel Xeon(R) CPU (E7-4830 v4) with 1056 GB of memory. Each tool was run with a single thread, and we recorded the wall clock time and the peak Resident Set Size (RSS) reported by the individual tools. We compared these with the output from GNU time 1.7 program and found no discrepancies.

### 2.3 Evaluations with mock community

We considered a real dataset consisting of Oxford Nanopore reads from the ZymoBIOMICS High Molecular Weight DNA Mock Microbial community (ZymoMC). We also used a simulated dataset of Oxford-like reads derived from the same genomes, but with a software-controlled error rate. The ZymoMC community consists of seven bacterial species – *Enterococcus faecalis*, *Listeria monocytogenes*, *Bacillus subtilis*, *Salmonella enterica*, *Escherichia coli*, *Staphylococcus aureus* and *Pseudomonas aeruginosa* – as well as *Saccharomyces cerevisiae* (yeast). As in prior studies [2, 4], we supposed that our goal was to deplete the bacterial reads, leading to proportionally more yeast reads sequenced.

#### 2.3.1 Assessing genomic versus pan-genomic indexes

We hypothesized that a pan-genome index – consisting of many related strains – would allow us to both (a) target a particular strain for depletion or enrichment when that specific strain is not present in the index, and (b) target a species as a whole by including many relevant strains or individuals from that species in the index. More specifically, we used the ZymoMC data and supposed that the seven bacterial strains were depletion targets (as in prior work [2, 4]). We assessed four strategies: (a) “One Genome w/o Zymo Mock Refs”: a single random strain from each of the seven bacterial species in ZymoMC, not matching the particular strain targeted for depletion, (b) “One Genome with Zymo Mock Refs”: the exact seven strains targeted for depletion, (c) “Pan-genome w/o Zymo Mock Refs”: all RefSeq strains for each bacterial species in ZymoMC, but excluding the depletion targets, and (d) “Pan-genome with Zymo Mock Refs”: all RefSeq strains for each bacterial species in ZymoMC including the depletion targets.

Table 1 shows that using an index containing many strains but excluding the specific depletion target yields a similar F1-score (99.7% for SPUMONI and minimap2) compared to when we use an index consisting only of the depletion target (99.1% for minimap2, 99.8% for SPUMONI). The F1 score remained unchanged when the pan-genome index was used.

**Table 1:**
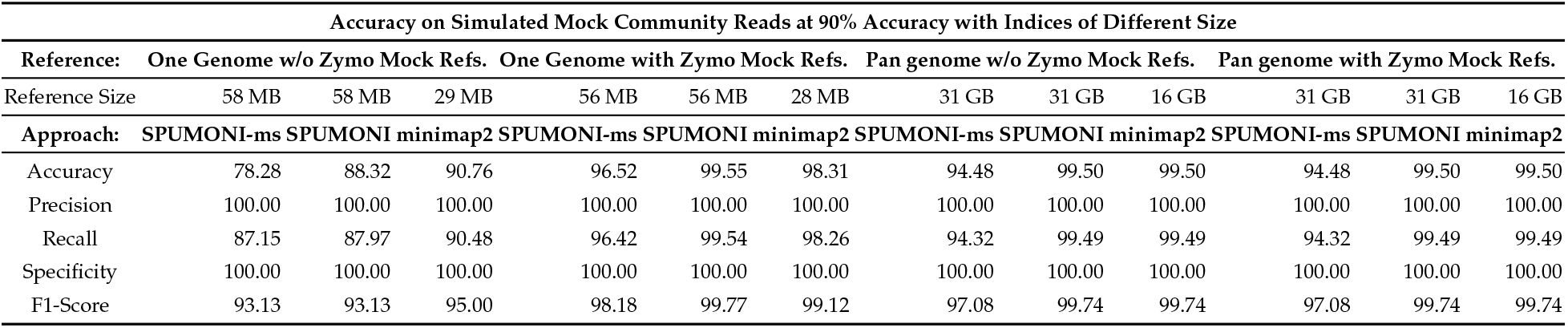
Assessing SPUMONI and minimap2 using both genomic and pan-genomic indexes. “SPUMONI” refers to the mode that uses PMLs, while SPUMONI-ms refers to the mode that uses matching statistics instead.

We conclude that a pan-genome index is a flexible tool for targeted sequencing, enabling targeting both at higher taxonomic levels, and in situations where the particular target strain has not been assembled or is unknown. In subsequent experiments, we continued to assess both a single-strain index (“One Genome w/o Zymo Mock Refs”) and a pan-genomic index (“Pan-Genome w/o Zymo Mock Refs”), focusing only on the indexes that exclude the target strain.

#### 2.3.2 Simulated Mock Community: Accuracy & Efficiency

To assess these methods in the presence of sequencing error, we used PBSIM2 [7] to simulate Oxford-Nanopore-like reads (R9.4 chemistry) from ZymoMC references at varying levels of mean read accuracy (%): 85, 90, 95, and 98. We again supposed that our goal was to eject reads from the seven bacterial strains so as to obtain proportionally more reads from yeast. The proportions of reads simulated from each genome was set to mimic those from the UNCALLED study [4] (Supplemental Figure 1). Figure 1 shows that as the error rate decreases, the distribution of matching statistics from the positive index gains a heavier right tail; that is, the half-maximal exact matches become longer since they are interrupted less often by sequencing errors.

**Figure 1:**
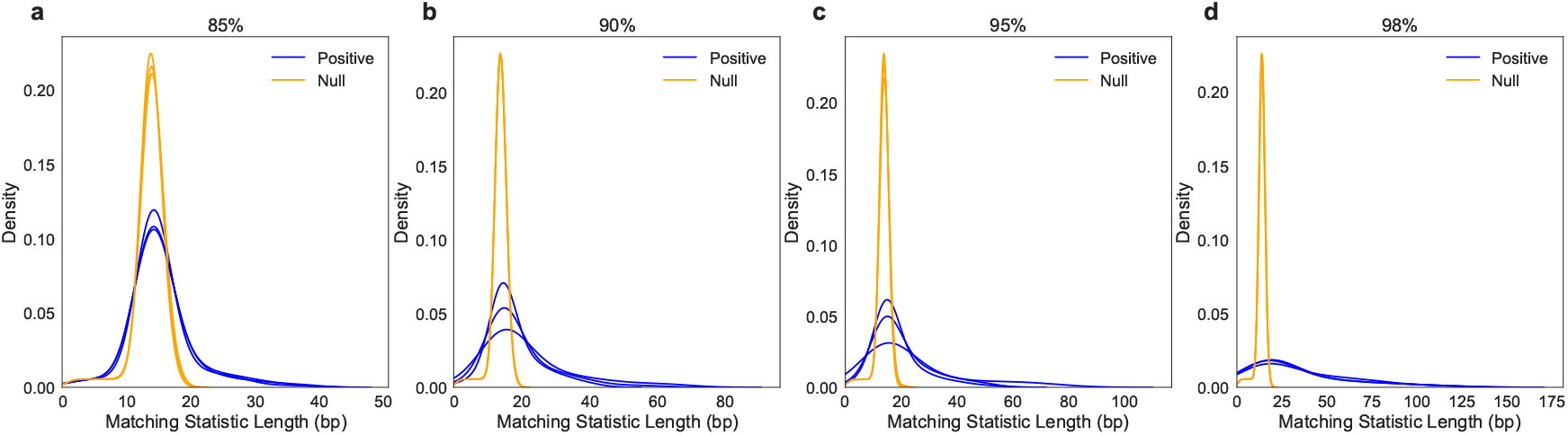
Distribution of matching statistics from positive and null indexes on simulated ZymoMC reads at accuracies of (a) 85%, (b) 90%, (c) 95%, and (d) 98%. Each plot contains the density curves for the first 720 bases (~ 1.6 seconds) for three randomly chosen simulated *Escherichia coli* reads.

We next compared SPUMONI to a minimap2-based approach, using the reads’ true simulated point of origin as the ground truth. As seen in Table 2, SPUMONI’s F1 score – and several related measures – increase as read accuracy increases. For reads at 90% accuracy and above, SPUMONI’s pan-genome index achieved ≥ 99.7% F1, which was comparable to and sometimes greater than minimap2’s pan-genome F1 scores. For both tools, the pan-genomic index substantially increased the F1 score, which is consistent with our results in Section 2.3.1.

**Table 2:** Comparing SPUMONI and minimap2 across various metrics on simulated ZymoMC reads of varying levels of accuracy. *The reported index size for SPUMONI-ms and SPUMONI includes only the positive index and not the null index, since the null index can be used offline and deleted prior to the analysis.

| Accuracy, Throughput and Index Size on Simulated Mock Community Reads at Various Levels of Accuracy |  |  |  |  |  |  |  |  |  |  |  |  |
| --- | --- | --- | --- | --- | --- | --- | --- | --- | --- | --- | --- | --- |
| Read Accuracy (%) | 85 |  |  |  |  |  | 90 |  |  |  |  |  |
| Reference: | One Genome Ref |  |  | Pan-genome Ref |  |  | One Genome Ref |  |  | Pan-genome Ref |  |  |
| Reference Size | 56 MB | 56 MB | 28 MB | 31 GB | 31 GB | 16 GB | 56 MB | 56 MB | 28 MB | 31 GB | 31 GB | 16 GB |
| Approach: | SPUMONI-ms | SPUMONI | minimap2 | SPUMONI-ms | SPUMONI | minimap2 | SPUMONI-ms | SPUMONI | minimap2 | SPUMONI-ms | SPUMONI | minimap2 |
| Accuracy | 56.21 | 83.35 | 87.53 | 70.08 | 95.43 | 99.16 | 78.28 | 88.32 | 90.76 | 94.48 | 99.50 | 99.50 |
| Precision | 100.00 | 100.00 | 100.00 | 100.00 | 100.00 | 100.00 | 100.00 | 100.00 | 100.00 | 100.00 | 100.00 | 100.00 |
| Recall | 54.89 | 82.85 | 87.15 | 69.18 | 95.29 | 99.13 | 87.15 | 87.97 | 90.48 | 94.32 | 99.49 | 99.49 |
| Specificity | 100.00 | 100.00 | 100.00 | 100.00 | 100.00 | 100.00 | 100.00 | 100.00 | 100.00 | 100.00 | 100.00 | 100.00 |
| F1-Score | 70.88 | 90.62 | 93.13 | 81.78 | 97.59 | 99.56 | 93.13 | 93.13 | 95.00 | 97.08 | 99.74 | 99.74 |
| Peak RSS (GB) | 0.63 | 0.08 | 0.17 | 6.24 | 1.90 | 8.07 | 0.63 | 0.08 | 0.17 | 6.24 | 1.90 | 8.07 |
| Index Size (GB)* | 0.68 | 0.09 | 0.10 | 6.20 | 1.90 | 31.00 | 0.68 | 0.09 | 0.10 | 6.20 | 1.90 | 31.00 |
| Throughput (bp/sec) | 134,690 | 614,665 | 398,415 | 28,731 | 111,813 | 6,441 | 177,572 | 709,018 | 409,104 | 33,829 | 125,914 | 6,617 |
| Read Accuracy (%) | 95 |  |  |  |  |  | 98 |  |  |  |  |  |
| Reference: | One Genome Ref |  |  | Pan-genome Ref |  |  | One Genome Ref |  |  | Pan-genome Ref |  |  |
| Reference Size | 56 MB | 56 MB | 28 MB | 31 GB | 31 GB | 16 GB | 56 MB | 56 MB | 28 MB | 31 GB | 31 GB | 16 GB |
| Approach: | SPUMONI-ms | SPUMONI | minimap2 | SPUMONI-ms | SPUMONI | minimap2 | SPUMONI-ms | SPUMONI | minimap2 | SPUMONI-ms | SPUMONI | minimap2 |
| Accuracy | 86.48 | 91.00 | 90.86 | 99.60 | 99.65 | 99.40 | 89.76 | 92.20 | 91.30 | 99.65 | 99.60 | 99.50 |
| Precision | 100.00 | 99.94 | 100.00 | 100.00 | 99.95 | 100.00 | 100.00 | 100.00 | 100.00 | 100.00 | 99.95 | 100.00 |
| Recall | 86.07 | 90.78 | 90.58 | 99.59 | 99.69 | 99.39 | 89.45 | 91.96 | 91.04 | 99.64 | 99.64 | 99.49 |
| Specificity | 100.00 | 98.31 | 100.00 | 100.00 | 98.31 | 100.00 | 100.00 | 100.00 | 100.00 | 100.00 | 98.31 | 100.00 |
| F1-Score | 92.52 | 95.14 | 95.06 | 99.80 | 99.82 | 99.69 | 94.43 | 95.81 | 95.31 | 99.82 | 99.80 | 99.74 |
| Peak RSS (GB) | 0.63 | 0.08 | 0.17 | 6.24 | 1.90 | 8.07 | 0.63 | 0.08 | 0.17 | 6.24 | 1.90 | 8.07 |
| Index Size (GB)* | 0.68 | 0.09 | 0.10 | 6.20 | 1.90 | 31.00 | 0.68 | 0.09 | 0.10 | 6.20 | 1.90 | 31.00 |
| Throughput (bp/sec) | 209,163 | 790,409 | 425,320 | 37,130 | 135,463 | 6,672 | 235,712 | 928,459 | 479,523 | 38,556 | 129,574 | 6,259 |

Considering throughput as measured in base pairs processed per second (bp/sec), SPUMONI is on average about 19.3 times faster than minimap2 when using the pan-genomic index, and about 1.8 times faster using the genomic index, and this is visualized in Supplemental Figure 2. Further, SPUMONI’s pan-genomic index is about 16 times smaller than minimap2’s, and SPUMONI’s peak memory footprint is about 4 times lower.

#### 2.3.3 Real Mock Community: Accuracy & Efficiency

Next, we applied our method to real nanopore reads from ZymoMC, obtained from SRA accession SRX7711546 [4]. When we plotted the distribution of matching statistics obtained from reads from different species, we observed that the distributions were quite distinct for the bacterial reads, but overlapping for the yeast (Supplemental Figure 3). This visualization shows how SPUMONI can distinguish between reads that it will try to eject and reads that it will let pass through the pore, and this difference can be statistically shown by differences in the KS-stat between the bacterial reads and the yeast reads (Supplemental Figure 4).

We compared SPUMONI to minimap2, this time using a separately-obtained minimap2 mapping as the gold standard. Specifically, we used minimap2 to map a suffix of the read, omitting the first 720 bases. To ensure the reads were long enough to enable an accurate mapping, we first filtered out reads that were shorter than 4,000 bp. We also trimmed the first 720 bases from each read before performing the gold-standard alignment, since these bases are used for classification later. Gold-standard labels were given only to reads that minimap2 could uniquely map to a ZymoMC reference with a MAPQ of ≥ 30. For reads that had at least one secondary alignment, we required that the ratio of the secondary alignment’s MAPQ to the primary alignment’s MAPQ was ≤ 0.60.

Results in Table 3 show that SPUMONI achieved similar F1 score as minimap2. For the genomic (“One Genome”) reference SPUMONI achieved 92.79% F1 score while minimap2 achieved 93.42% F1 score. Both tools achieved 100% precision and specificity in this case. For the pan-genomic (“Pan-genome”) reference, SPUMONI achieved 97.94% F1 score while minimap achieved 98.73%. In this case, SPUMONI achieved 100% precision and specificity whereas minimap2 achieved 99.96% precision and 96.97% specificity.

**Table 3:** Comparing SPUMONI and minimap2 across various metrics on Real ZymoMC Reads. *The index sizes for SPUMONI-ms and SPUMONI are only for the positive index, since the null index can be used offline and removed.

| Accuracy, Throughput and Index Size on Real Mock Community Reads |  |  |  |  |  |  |
| --- | --- | --- | --- | --- | --- | --- |
| Reference: | One Genome Ref |  |  | Pan-genome Ref |  |  |
| Reference Size | 56 MB | 56 MB | 28 MB | 31 GB | 31 GB | 16 GB |
| Approach: | SPUMONI-ms | SPUMONI | minimap2 | SPUMONI-ms | SPUMONI | minimap2 |
| Accuracy | 81.64 | 86.72 | 87.82 | 94.62 | 96.02 | 97.52 |
| Precision | 100.00 | 100.00 | 100.00 | 100.00 | 100.00 | 99.96 |
| Recall | 81.39 | 86.54 | 87.66 | 94.55 | 95.97 | 97.53 |
| Specificity | 100.00 | 100.00 | 100.00 | 100.00 | 100.00 | 96.97 |
| F1-Score | 89.74 | 92.79 | 93.42 | 97.20 | 97.94 | 98.73 |
| Peak RSS (GB) | 0.63 | 0.08 | 0.17 | 6.24 | 1.90 | 8.07 |
| Index Size (GB)* | 0.68 | 0.09 | 0.10 | 6.20 | 1.90 | 31.00 |
| Throughput (bp/sec) | 252,974 | 901,609 | 851,869 | 64,384 | 185,618 | 15,570 |

When using the pan-genome reference, SPUMONI achieved a throughput about 11.9 times higher than that of minimap2. While when using the genomic index, SPUMONI achieved slightly higher throughput compared to minimap2 (902 kbp/sec versus 852 kpb/sec). When measuring peak Resident Set Size (RSS), we observed that SPUMONI’s memory footprint was about 1/4th that of minimap2, and that its index was about 16 times smaller.

### 2.4 Human Microbiome

Finally, we assessed our method on a human microbiome sequencing scenario with the goal of ejecting reads from the human host to enrich for any microbial species present. We constructed a dataset consisting of a mixture of real reads from a recent human microbiome study that used Oxford Nanopore sequencing [8], as well as a set of simulated human nanopore-like reads with a mean read accuracy of 90%. Likely-human reads were already filtered out of the former dataset; therefore, we assumed that the only human reads in the final read set are the simulated ones. Since a human genome assembly is on the order of 3 billion nucleotides, an index containing one or more human assemblies presents a significantly larger but relevant challenge.

When we visualized the distribution of matching statistics for reads from different species (Figure 2), we saw the simulated human reads appeared to match the positive index (evidenced by the blue densities’ thicker right tails) while reads from the microbiome study did not (indicated by the similarity of positive and null distributions).

**Figure 2:**
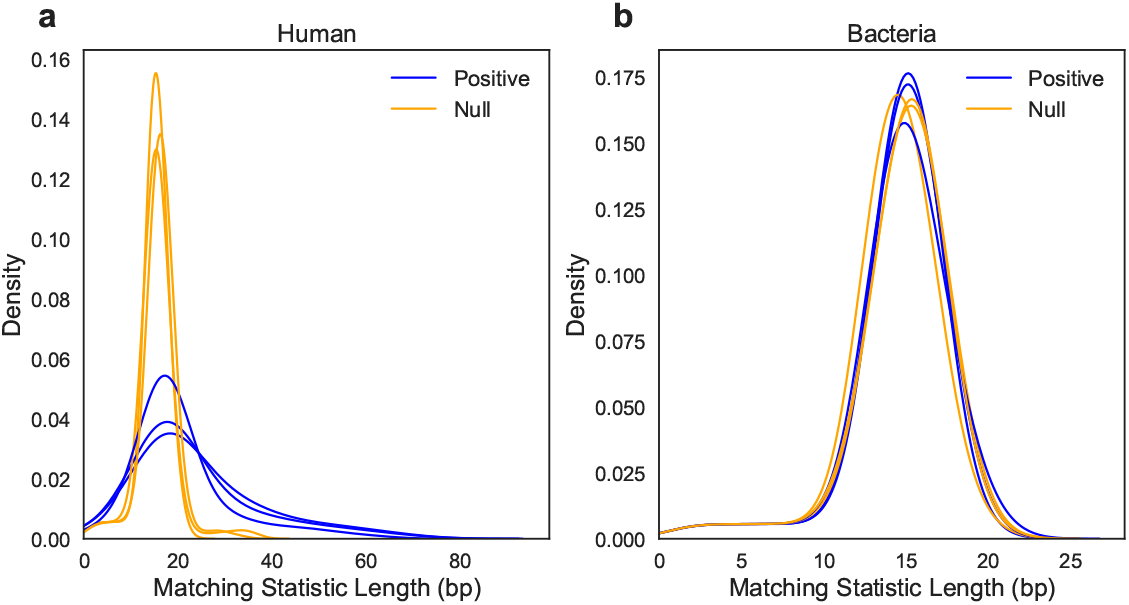
Distribution of matching statistics across three randomly chosen reads from (a) the human simulation and (b) the microbiome study [8]. A single curve represents the first 720 bases (~ 1.6 Read Until seconds) of a read.

We evaluated SPUMONI and minimap2 on this dataset using two different indexes: (a) an index consisting only of the Telomere-to-Telomere Consortium (“T2T”) CHM13 v1.0 assembly [9], and (b) an index consisting of the T2T assembly together with the Ashkenazi [10] and GRCh38 [11] assemblies. Indexing multiple human genomes allows us to achieve similar benefits as we did for the mock-community pan-genomes, i.e. coverage of a wider range of genetic variation, particularly structural variation. It also helps to reduce reference bias, which in our case would manifest as a tendency to find shorter matches in genomic regions with non-reference alleles.

When using the single-genome index, SPUMONI achieved somewhat higher F1 score (96.97%) compared to minimap2 (95.17%), and lower throughput (24.5 versus 35.7 kpb/sec). When using the 3-genome index, minimap2 achieved higher F1 score (99.17%) than SPUMONI (97.08%) but SPUMONI had higher throughput (27.0 versus 13.5 kbp/src), which is shown in Table 4. As the reference became more repetitive – moving from one to 3 genomes – SPUMONI gained an index-size advantage, using 18 GB versus minimap2’s 21 GB for the 3-genome index. While this comparison between SPUMONI and minimap2 is close, we expect that as we are able to index and align to more human references simultaneously — e.g. as more assemblies from the Human Pan-Genome Reference Consortium [12] and similar projects emerge — SPUMONI is well positioned for sublinear index growth and a greater throughput advantage. For instance, the *r*-index underlying SPUMONI was previously shown to be able to index up to 10 human genomes with sublinear growth in the index size [13].

**Table 4:** Comparing SPUMONI and minimap2 on various metrics when processing the human microbiome reads. *The index sizes for SPUMONI-ms and SPUMONI are only for the positive index, since the null index can be used offline and removed.

| Accuracy, Throughput and Index Size on Human Microbiome Reads |  |  |  |  |  |  |
| --- | --- | --- | --- | --- | --- | --- |
| Reference: | One Human Genome |  |  | Three Human Genomes |  |  |
| Reference Size | 5.8 GB | 5.8 GB | 2.9 GB | 18.0 GB | 18.0 GB | 9.0 GB |
| Approach: | SPUMONI-ms | SPUMONI | minimap2 | SPUMONI-ms | SPUMONI | minimap2 |
| Accuracy | 98.66 | 99.42 | 99.10 | 98.64 | 99.44 | 99.84 |
| Precision | 95.46 | 98.93 | 100.00 | 95.06 | 98.73 | 100.00 |
| Recall | 90.57 | 95.08 | 90.78 | 90.78 | 95.49 | 98.36 |
| Specificity | 99.54 | 99.89 | 100.00 | 99.49 | 99.87 | 100.00 |
| F1-Score | 92.96 | 96.97 | 95.17 | 92.87 | 97.08 | 99.17 |
| Peak RSS (GB) | 57.29 | 15.23 | 7.85 | 62.80 | 18.06 | 9.70 |
| Index Size (GB)* | 57.90 | 15.00 | 6.90 | 62.60 | 18.00 | 21.00 |
| Throughput (bp/sec) | 7,518 | 24,476 | 35,742 | 6,860 | 27,024 | 13,459 |

## 3 Discussion

SPUMONI is a streaming algorithm for targeted nanopore sequencing that uses matching statistics (and “pseudo matching lengths”) to classify reads in real-time. SPUMONI’s data structures – the *r*-index and MONI thresholds – allow it to handle repetitive pan-genome indexes more efficiently than competing approaches. SPUMONI’s memory efficiency combines well with the flexibility afforded by nanopore sequencing, allowing SPUMONI to run on more portable hardware, like that associated with MinION and Flongle instruments. The ability to include a wide array of strains in a single index makes SPUMONI attractive for metagenomics applications where targets may not have already been cultured, assembled and deposited in a resource like Refseq. As nanopore sequencing continues to improve, both base-calling accuracy and per-instruction throughput will likely improve. SPUMONI is well positioned for these trends, since it delivers its most advantageous combinations of speed and F1 score at higher base calling accuracy.

SPUMONI operates on batches of already-called bases. In practice, the Read Until API delivers data in the form of raw current that must be base-called first. Since nanopore base-callers have been steadily improving, it is possible that base calling will be integrated into on-board components of nanopore sequencers. Until then, users must run a separate base caller upstream of SPUMONI, as also required by Readfish [2]. That said, the fact that SPUMONI’s analysis is at the level of bases allows it to target other classification problems, like metagenomics classificaton.

While SPUMONI’s null index currently consists of the reverse of the sequences used in the positive index, this notion of “null” might be insufficient in some scenarios. For example, if there is substantial sequence similarity between depletion-target reads and reads that should not be targeted for depletion – e.g. due to conserved genes between species – the positive matching statistics within those sequences will be longer than what is expected by random chance for depletion-target reads. In these cases, we may need to augment the null model, perhaps by including the conserved sequences (not their reverses) in the null index.

Currently, we use the same KS-stat threshold for all the experiments which was optimized to perform well on real nanopore datasets. However, we expect that the optimal threshold will also be a function of the read accuracy and the reference used. In future work, we will investigate whether a simulation could be used to model the sequencing run to determine a threshold that is more tailored to a particular experiment.

Finally, we observed that SPUMONI can compress reads as it processes them: we can simply output each PML followed by the character in the read that did not match the corresponding character in the BWT; to decompress the read, we recover the characters that matched (and caused the PML to increment) using LF steps until we reach the mismatch character, at which point we jump to the previous or next occurrence of that character in the BWT, as we did while compressing the read. Since the compressing ratio of this scheme improves with larger PMLs, we may be able to use that compression ratio as an aggregate statistic when deciding whether to eject a read. Finally, we note that in some sense this compression scheme works by predicting the characters in the reads and recording explicitly those characters it predicts incorrectly.

## 4 Methods

### 4.1 Matching statistics with r-index

Given a text *T*[1..*n*] of length *n*, the Burrows-Wheeler Transform (BWT) [14] is a reversible permutation of the *T* such that the character in position *i* is the character preceding the *i*-th lexicographic sorted suffix of *T*. We use *r* to denote the number of maximal equal-letter runs of the BWT. The *r*-index [13] is a self-index which stores a run-length encoded BWT, i.e., each run is encoded as a character together with the run length.

Given a text *T*[1..*n*] of length *n* and a pattern *P*[1..*m*] of length *m*, the *matching statistics* of *P* against *T* are defined as an array *MS*[1..*m*] of length *m*, where each position *MS*[*i*] stores the length of the longest prefix of *P*[*i*..*m*] that occurs in *T*. Bannai et al. [15] introduced the *thresholds* which are *O*(*r*) positions in the BWT marking a minimum of the longest common prefix array, between two equal letter runs. They also proposed a two-pass algorithm to compute matching statistics using use these thresholds and the *r*-index. In the first pass, the algorithm steps backward along the pattern *P*. When it can, the algorithm uses the LF mapping to extend the match by one character. Where this is not possible, we “jump” either forward or back in the BWT to a position where the match can be extended. Whether we jump forward or back depends on which direction gives the longer common prefix with the match so far, which in turn is determined by the threshold’s location. In the second pass, the algorithm uses a random-access data structure built over *T* to compute the lengths of the matching statistics.

Rossi et al. [6] with MONI showed how to efficiently compute the thresholds for highly repetitive texts, and implemented the matching statistics algorithm. A MONI index consists of four main components, the run-length encoded BWT, suffix-array samples taken at run boundaries, the thresholds, and a grammar [16, 17] that provides random access to *T*. These data structures allow computation of matching statistics in *O*(*m* log *n*) time, and take *O*(*r* + *g*) space where *g* is the size of a given straight-line program (SLP) for *T*.

### 4.2 Pseudo matching lengths

SPUMONI modifies MONI by removing the second pass. As SPUMONI performs a backward LF-mapping search, it increments a length variable whenever the BWT character encountered matches the next character in *P*. If the character fails to match, the length variable is reset to 0 and we “jump” in the BWT as usual. The value of the length variable at each step gives the sequence of *pseudo matching lengths* (PMLs); these differ from matching statistics since we have ignored the possibility that a BWT jump can correspond to an extension of the current half-maximal match. PMLs will consistently be shorter than the true MSs. But long MSs – long enough to narrow the BWT range to the point where random matches are excluded – will generally yield long PMLs. Since the longest MSs are the ones with the most power to discriminate target from non-target, we expect, and our results confirm, that PMLs are similarly useful for classification.

This simplification obviates the need to store either the SA samples or the random-access grammar for *T*; those were used only in MONI’s second loop. Hence, a SPUMONI index consists only of the run-length encoded BWT and thresholds. This leads to improvements in the time and space complexity, where pseudo matching lengths can be computed in *O*(*m* log log *n*) time and take *O*(*r*) space in worst case. Pseudocode highlighting differences between MONI and SPUMONI – and between MSs and PMLs – is given in Supplemental Figure 5.

### 4.3 Positive and Null Indexes

In our approach, we generated matching statistics of the read with respect to both a positive and null index. The positive index consisted of both the forward and reverse complement of the sequence that we wanted to target, whether that be for depletion or enrichment. The null index simply consisted of the reverse of the positive index sequence, and the matching statistics generated with respect to the null index were meant to represent matching statistics you would get against random sequence. This would allow us to compare the distribution of matching statistic lengths with respect to the positive index to a baseline distribution, and if we see a clear difference, it is probably due to the read matching significantly to sequence in the positive index.

### 4.4 Statistical Test

To decide whether the positive and null distributions of matching statistics are different, we used the Kolmogorov-Smirnov test (KS-test), which compares the distributions’ cumulative distribution functions (CDFs). We found that a Kolmogorov-Smirnov statistic (KS-stat) 0.25 and 0.10 for matching statistics and pseudo matching lengths respectively worked well across different nanopore datasets. We applied the KS-test to non-overlapping regions of 90 bp which allows to us to compute the KS-stat as the Read Until API delivers new batches of data without having to revisit and use earlier batches of data in the computation.

In addition, prior to feeding in the matching statistics from the non-overlapping regions into the KS-test, we applied a transformation function to the data. The function consisted of taking each matching statistic length and subtracting the mean of the null distribution, and replacing its value with 1 if it was less than 1. The intuition behind this function is that it compresses all of matching statistic lengths that are near-random length into a matching statistic length of 1. This improves the accuracy using the KS-test because the KS-test is based on distances between CDFs so this transformation will tend to increase the KS-stat when the distributions are truly different.

Finally, in order to make a decision on the read-level for whether the read should be classified as matching sequence in the positive index or not. We gather all the KS-stats from the non-overlapping regions and see if a simple majority of them are above the threshold.

### 4.5 Datasets

For the “Pan-genome Reference” collection in Section 2.3.1, we used all available genomes for each bacterial species of the ZymoMC in the RefSeq Database. Accession numbers for the bacterial genomes can be downloaded at https://benlangmead.github.io/aws-indexes/ spumoni, and the number of genomes in each species can be found in Supplemental Table S1. For the human assemblies in Section 2.4, we used the Telomere-to-Telomere Consortium CHM13 v1.0 assembly [9], the Ashkenazi assembly [10], and GRCh38 [11].

For Section 2.3.3, we used the reads present in the SRA Project under Accession Number SRX7711546 [4]. The reads were pre-filtered by aligning the entire reads using minimap2 [3] and keeping the reads that mapped uniquely to the ZymoMC references with MAPQ of at least 30. We also kept the reads where the ratio of the MAPQ of their secondary and primary alignments was no more than 0.60. For read sets used in Section 2.4, the human reads were simulated from the Telomere-to-Telemere Consortium CHM13 v1.0 assembly [9] at a mean read accuracy of 90% using PBSIM2 [7] and its model for the R9.4 chemistry. The bacterial microbiome reads were obtained SRA Accession SRX6602475 [8].

SPUMONI indexes used for each experiment can be obtained from: https://benlangmead.github.io/aws-indexes/spumoni.

## 5 Acknowledgments

Part of this research project was conducted using computational resources at the Maryland Advanced Research Computing Center (MARCC). SPUMONI indexes are made freely available on Amazon Web Services thanks to the AWS Public Dataset Program.

