## Supplemental Figure 1 for "Pan-genomic Matching Statistics for Targeted Nanopore Sequencing"

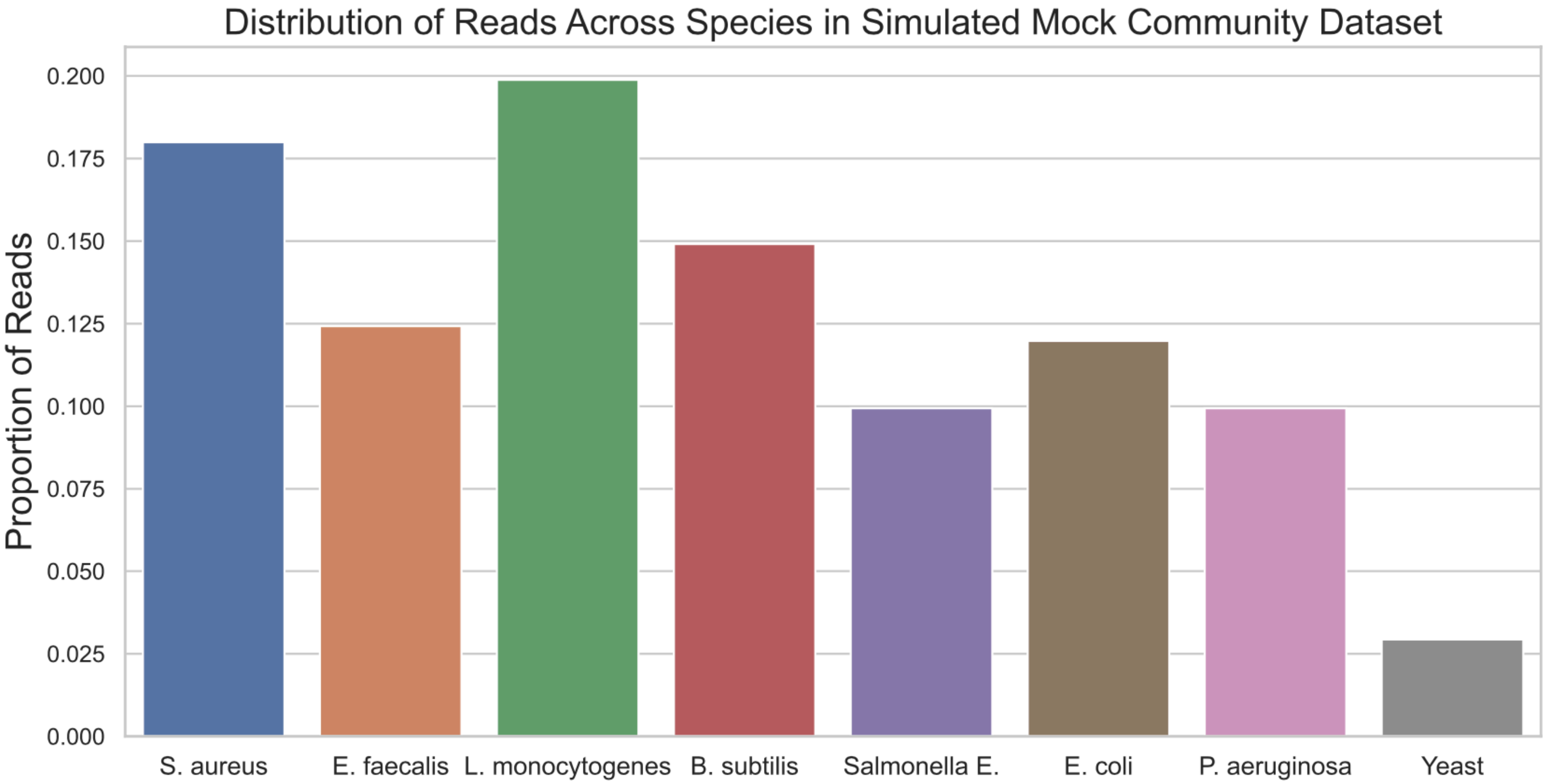


Supplementary Figure 1: Proportion of reads from each species in mock community in the simulated mock community dataset.
