## Supplemental Figure 2 for "Pan-genomic Matching Statistics for Targeted Nanopore Sequencing"

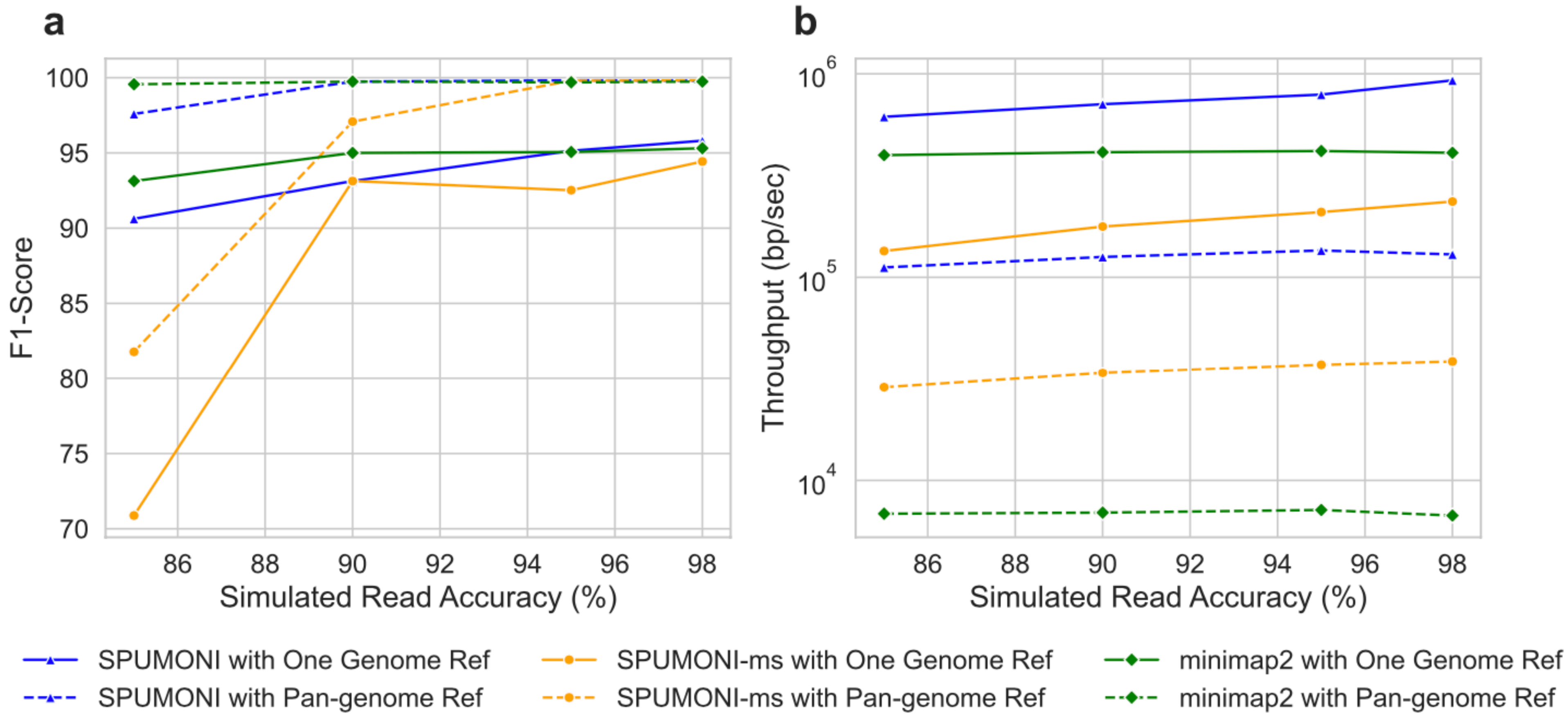


Supplementary Figure 2: Visualization of (a) $F_1$-score, (b) throughput with varying simulated base-call accuracies.
