## Supplemental Figure 3 for "Pan-genomic Matching Statistics for Targeted Nanopore Sequencing"

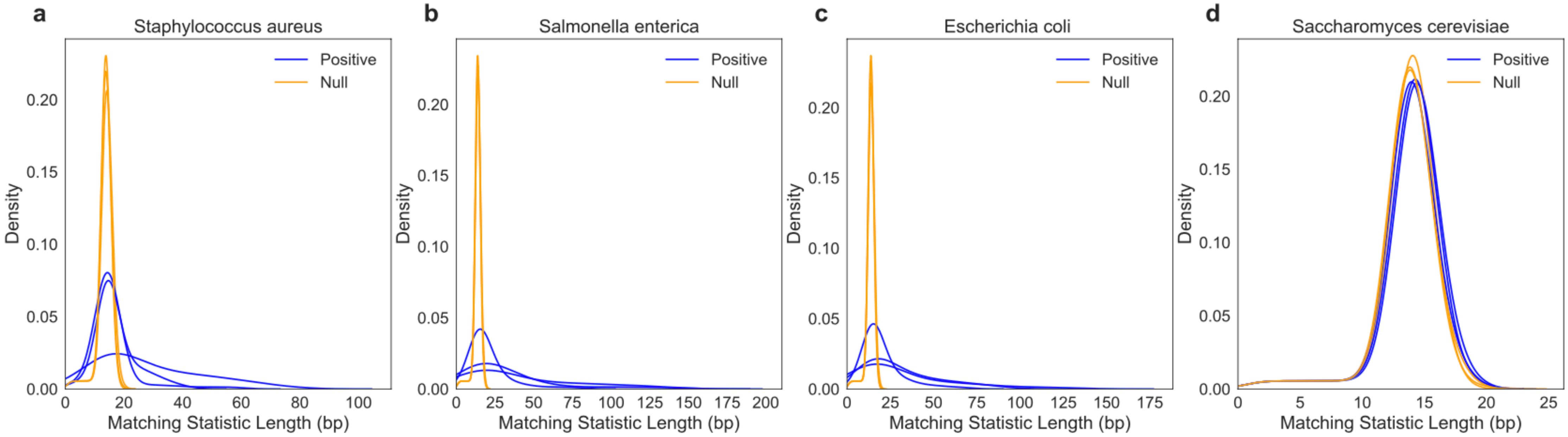


Supplementary Figure 3: Distribution of matching statistics against positive and null indexes for three randomly chosen reads of (a) *Staphylococcus aureus*, (b) *Salmonella enterica*, (c) *Escherichia coli*, and (d) *Saccharomyces cerevisiae*. Each density curve is based on the first 720 bases ($\sim$ 1.6 seconds) of ReadUntil data for each read.
