## Supplemental Figure 4 for "Pan-genomic Matching Statistics for Targeted Nanopore Sequencing"

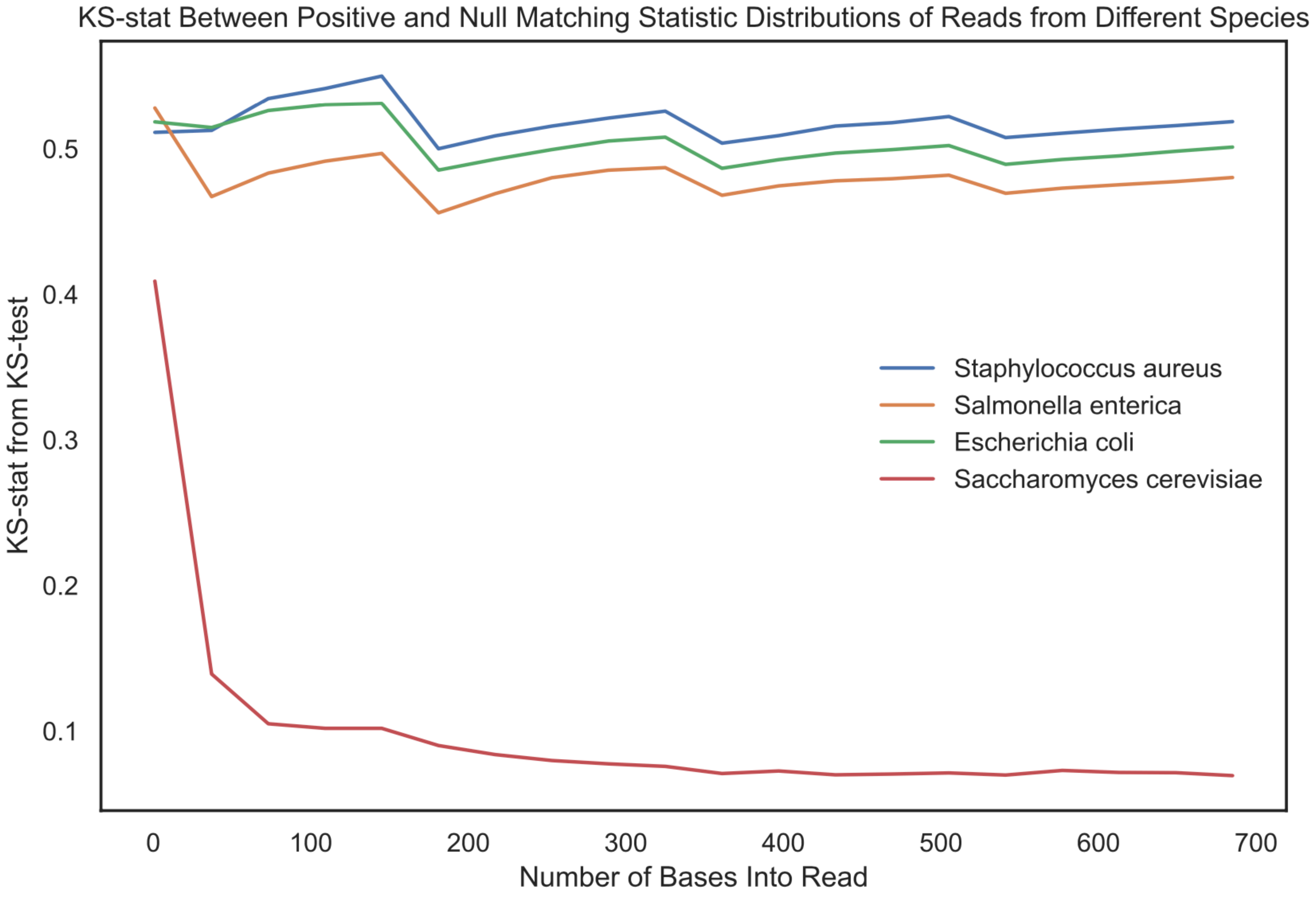


Supplementary Figure 4: Each line represents the average of the Kolomgorov-Smirnov statistic (KS-stat) across all reads from that respective species in the mock community. The figure shows that for bacterial species (*Staphylococcus aureus*, *Salmonella enterica*, and *Escherichia coli*) the KS-stat is relatively larger at $\sim 0.5$ since the reads are matching to sequence in the positive index. While for the yeast reads, the KS-stat is small at $\sim 0.1$ since the distribution of positive and null matching statistics are quite similar to each other.
