## Supplemental Figure 5 for "Pan-genomic Matching Statistics for Targeted Nanopore Sequencing"

**Algorithm 1** Matching Statistics

```
given text T and a pattern p
//first pass: get pointers
j ← 1, pos ← SA[j]
for i ← p.len to 1 do
    if p[i] ≠ BWT[j] then
        if j < Thr(j, p[i]) then
            j ← BWT.pred(j, p[i])
        else
            j ← BWT.succ(j, p[i])
        pos ← SA[j] − 1
    else
        pos ← pos − 1
    pointers[i] ← pos
    j ← LF(j)
//second pass: get lengths
ℓ ← 0
for i ← 1 to pointers.len do
    pos ← pointers[i] + ℓ, j ← i + ℓ
    while j < p.len and p[j] = T[pos] do
        ℓ ← ℓ + 1
        pos ← pos + 1, j ← j + 1
    ms[i] ← ℓ
    ℓ ← max(ℓ − 1, 0)
Return ms
```

**Algorithm 2** Pseudo Matching Lengths

```
given text T and a pattern p
//first pass: get pointers and pmls
j ← 1, ℓ ← 0
for i ← p.len to 1 do
    if p[i] ≠ BWT[j] then
        if j < Thr(j, p[i]) then
            j ← BWT.pred(j, p[i])
        else
            j ← BWT.succ(j, p[i])
        ℓ ← 0
    else
        ℓ ← ℓ + 1
    pml[i] ← ℓ
    j ← LF(j)
Return pml
```

Supplementary Figure 5: Matching statistics (Algorithm 1) and pseudo matching lengths (Algorithm 2) computation using the thresholds. Given an array $a$, $a.len$ refers to the length of the array. Given a position $j$ in the BWT, $\mathrm{LF}(j)$ is the LF-mapping, SA is the suffix array sampled at run boundaries, $\mathrm{BWT.pred}(j, c)$ is the position in the BWT of the first character $c$ preceding position $j$, $\mathrm{BWT.succ}(j, c)$ is the position in the BWT of the first character $c$ following position $j$, $\mathrm{Thr}(j, c)$ is the position of the threshold for the character $c$ between the run of $c$ preceding and following position $j$ in the BWT. To avoid overloading the notation, we consider the variable $pos$ to be $(pos \bmod T.len) + 1$. Furthermore, note that if $\mathrm{BWT.pred}(j, c)$, the threshold value is guaranteed to be smaller than or equals to $j$, i.e., $\mathrm{Thr}(j, c) = 0$. Similarly, if $\mathrm{BWT.succ}(j, c)$, the threshold value is guaranteed to be greater than $j$, i.e., $\mathrm{Thr}(j, c) = T.len + 1$.
