## Supplemental Table S1 for "Pan-genomic Matching Statistics for Targeted Nanopore Sequencing"

| **Number of Genomes Included in Each Mock Community Index** | | |
|---|---|---|
| **Species\Reference:** | One Genome Ref. | Pan-genome Ref. |
| Staphylococcus aureus | 1 | 574 |
| Enterococcus faecalis | 1 | 49 |
| Listeria monocytogenes | 1 | 225 |
| Bacillus subtitles | 1 | 165 |
| Salmonella enterica | 1 | 880 |
| Escherichia coli | 1 | 1370 |
| Pseudomonas aeruginosa | 1 | 274 |
| Saccharomyces cerevisiae | 1 | 1 |

Table S1: Number of genomes for each species in the different references used for the mock community experiment.
